# Radtools: R utilities for smooth navigation of medical image data

**DOI:** 10.1101/471706

**Authors:** Pamela H Russell, Debashis Ghosh

**Affiliations:** Department of Biostatistics and Informatics, Colorado School of Public Health, Aurora, CO, USA

**Keywords:** Medical imaging, DICOM, NIfTI, R package

## Abstract

The radiology community has adopted several widely used standards for medical image files, including the popular DICOM (Digital Imaging and Communication in Medicine) and NIfTI (Neuroimaging Informatics Technology Initiative) standards. These file formats include image intensities as well as potentially extensive metadata. The NIfTI standard specifies a particular set of header fields describing the image and minimal information about the scan. DICOM headers can include any of over 4,000 available metadata attributes spanning a variety of topics. NIfTI files contain all slices for an image series, while DICOM files capture single slices and image series are typically organized into a directory. Each DICOM file contains metadata for the image series as well as the individual image slice.

The programming environment R is popular for data analysis due to its free and open code, active ecosystem of tools and users, and excellent system of contributed packages. Currently, many published radiological image analyses are performed with proprietary software or custom unpublished scripts. However, R is increasing in popularity in this area due to several packages for processing and analysis of image files. While these R packages handle image import and processing, no existing package makes image metadata conveniently accessible. Extracting image metadata, combining across slices, and converting to useful formats can be prohibitively cumbersome, especially for DICOM files.

We present radtools, an R package for smooth navigation of medical image data. Radtools makes the problem of extracting image metadata trivially simple, providing simple functions to explore and return information in familiar R data structures. Radtools also facilitates extraction of image data and viewing of image slices. The package is freely available under the MIT license at https://github.com/pamelarussell/radtools and is easily installable from the Comprehensive R Archive Network (https://cran.r-project.org/package=radtools).

## Introduction

Medical image analysis often lies at the boundary of research and the clinic, presenting challenges in both domains. Institutional and privacy concerns can compete with the objective of open data for research purposes. In particular, it remains standard practice to perform analysis with proprietary software or unpublished scripts. Additionally, the majority of imaging studies do not make image data publically available due to patient privacy requirements. These complex challenges can present barriers for scientists working in the image analysis domain.

In recent years, a small but growing number of open source computational tools have been developed to process and analyze medical images, promoting sharing of code; some of the most widely adopted are described in (1–3). To address the issue of availability of public image data, our group previously developed TCIApathfinder (4), an open source R package to simplify access to the thousands of publicly available images in The Cancer Imaging Archive (5). Here, we present radtools, an open source R package that lowers barriers to image analysis by simplifying the extraction of image properties and complex header information. Although several excellent image processing and analysis packages exist for the R environment (2,6–9), none currently offers special functionality for convenient presentation of image metadata. Radtools implements a layer of processing to convert image metadata to familiar R data structures, eliminating the need for specialized knowledge and custom scripts to parse image files.

Radtools supports the two most common medical image formats, DICOM (Digital Imaging and Communication in Medicine) (10) and NIfTI-1 (Neuroimaging Informatics Technology Initiative) (11). The industry standard DICOM format combines a header and two-dimensional image data into one file, so that an image acquisition typically produces multiple DICOM files. DICOM header fields consist of a “tag” that identifies the attribute, followed by the attribute value. There is no fixed size for a DICOM header; any number of thousands of possible attributes may be included. Each DICOM file for an acquisition contains its own header; many attributes will be constant across image slices. NIfTI-1 format was developed primarily for multidimensional imaging data as an improvement over the previous ANALYZE format (12). NIfTI-1 combines header information and the entire multidimensional image acquisition into either a single file or two files (one header file and one image file). Unlike DICOM, NIfTI-1 specifies a particular set of required header attributes, and the header conforms to a fixed size with an option to add extended header information. Radtools provides simple functions to explore and return image properties and header data from both image formats in familiar R data structures. Radtools also facilitates extraction of image data and viewing of image slices.

## Methods

### Implementation

Radtools is provided as a package (extension to the language) for the programming language R. The package is hosted on the Comprehensive R Archive Network (CRAN), and can be installed into the user’s local R environment with the command ‘install.packages(“radtools”)’.

### Operation

The only system requirement is a working installation of R version ≥ 3.4.0. Radtools consists of a collection of functions that can be called within R scripts or interactively from the R console. Package operation is documented in a vignette that can be viewed on the GitHub page (https://github.com/pamelarussell/radtools), the CRAN page (https://cran.r-project.org/package=radtools), or from the R console with the command ‘browseVignettes(“radtools”)’. The package reference manual provides documentation of each individual function and is available on the CRAN page.

## Use cases

Radtools can extract image properties and header data from any valid DICOM or NIfTI-1 file. Image datasets are loaded with the ‘read_dicom’ and ‘read_niftil’ functions. Several generic functions extract attributes from either data type, including ‘img_dimensions’, ‘num_slices’, ‘header_fields’, which reports the set of header fields present, and ‘header_value’, which returns the value(s) of a particular attribute. Additionally, functions are provided to specifically address one format or the other. All header data present in a DICOM acquisition can be extracted into a matrix, where rows are attributes and columns are slices, with the ‘dicom_metadata_matrix’ function. As most DICOM headers contain numerous attributes and many of these are constant across all slices, the ‘dicom_constant_header_values’ function produces a named list of common attributes across slices. NIfTI-specific functions include ‘niftil_num_dim’, which returns the number of dimensions, and ‘niftil_header_values’, which returns a named list of all metadata attributes for the image.

The image itself can be extracted as a multidimensional matrix of intensities for either file format with ‘img_data_to_mat’. Image slices can be visualized with ‘view_slice’.

Finally, functions are provided to explore aspects of the DICOM standard itself. The functions ‘dicom_all_valid_header_tags’, ‘dicom_all_valid_header_names’, and ‘dicom_all_valid_header_keywords’ return complete lists of valid DICOM header attributes. The functions ‘dicom_search_header_names’ and ‘dicom_search_header_keywords’ return attributes matching a search term.

## Conclusions

Radtools fills a specific need in the existing ecosystem of R packages for image processing and analysis: namely, the need for smooth extraction of image metadata. The package will accelerate workflow development and provide researchers with easy access to attributes that they may not have otherwise considered using. The inclusion of the package on CRAN, along with clear documentation, make it trivially simple for R users to obtain and begin using radtools.

## Data availability

No data is associated with this article.

## Software availability

Radtools is available under the MIT license at https://doi.org/10.5281/zenodo.1477093. The development version is available at https://github.com/pamelarussell/radtools. Radtools is on CRAN at https://cran.r-project.org/package=radtools, and can be installed with the R command “install.packages(“radtools”).

## Author Contributions

P.R.: Conceptualization, Methodology, Software, Writing — Original Draft Preparation, Writing — Review & Editing

D.G.: Conceptualization, Funding Acquisition, Writing — Review & Editing

